# The paradox of Rapid Eye Movement sleep in the light of oscillatory activity and functional synchronization during phasic and tonic microstates

**DOI:** 10.1101/498212

**Authors:** Péter Simor, Gwen van Der Wijk, Ferenc Gombos, Ilona Kovács

**Author notes:** The authors contributed equally to this work (Péter Simor, Gwen van Der Wijk). Corresponding author: Peter Simor, Ph.D., Institute of Psychology, ELTE Eötvös Loránd University, Budapest, Hungary. 1064 Budapest, Izabella utca 46.

## Abstract

Rapid Eye Movement (REM) sleep is a peculiar neural state showing a combination of muscle atonia and intense cortical activity. REM sleep is usually considered as a unitary state in neuroscientific research; however, it is composed of two different microstates: phasic and tonic REM. These differ in awakening thresholds, sensory processing, and cortical oscillations. Nevertheless, studies examining cortical oscillations during REM microstates are scarce, and used low spatial sampling. Here, we analyzed the data of 18 healthy individuals assessed by high-density sleep EEG recordings. We systematically contrasted phasic and tonic REM periods in terms of topographical distribution, source localization, as well as local, global and long-range synchronization of frequency-specific cortical activity. Tonic periods showed relatively increased high alpha and beta power over frontocentral derivations. In addition, higher frequency components of beta power exhibited increased global synchronization during tonic compared to phasic states. In contrast, in phasic periods we found increased power and synchronization of low frequency oscillations coexisting with increased and synchronized gamma activity. Source localization revealed several multimodal, higher-order associative, as well as sensorimotor areas as potential sources of increased high alpha/beta power during tonic compared to phasic REM. Increased gamma power in phasic REM was attributed to medial prefrontal and right lateralized temporal areas associated with emotional processing. Our findings emphasize the heterogeneous nature of REM sleep, expressed in two microstates with remarkably different neural activity. Considering the microarchitecture of REM sleep may provide new insights into the mechanisms of REM sleep in health and disease.

**Significance Statement:** Rapid Eye movement (REM) is composed of the alternation of two markedly different microstates: phasic and tonic REM. Here we present the first high-resolution topographical study that systematically contrasted spectral power, neural synchronization and brain topography of spontaneous cortical oscillations across phasic and tonic microstates. Phasic REM was characterized by the combination of synchronized low frequency oscillations resembling deeper sleep stages and high frequency oscillations reflecting intense cortical activity. In contrast, tonic REM sleep showed increased oscillatory activity in the alpha and beta ranges that may reflect a shift towards increased environmental alertness resembling resting wakefulness. Exploring the neurophysiological aspects of REM microstates may shed new light on the mechanisms of REM sleep in healthy and pathological conditions.

## Introduction

Rapid Eye Movement (REM) sleep is a puzzling neural state characterized by bursts of rapid eye movements (REMs), low amplitude and mixed frequency electroencephalographic (EEG) oscillations, muscle atonia, irregular vegetative activity, and reduced thermoregulation (Siegel, 2005). Healthy human adults spend approximately 1.5 hours every night in REM sleep, that appear regularly after the ascending phases of non-REM (NREM) sleep and lasts around 20 minutes depending on circadian, homeostatic and other state factors (Carskadon and Dement, 2005). These periods bear some resemblance to wakefulness in terms of increased cortical (Muzur et al., 2002; Miyauchi et al., 2009) and neural network activity (Cantero et al., 2004; Massimini et al., 2010) that gives rise to peculiar conscious (oneiric) experiences. The function of REM sleep has been linked to a wide range of neural and cognitive phenomena such as cortical maturation (Blumberg et al., 2013), synaptic pruning (Li et al., 2017), emotional regulation (Walker and van der Helm, 2009), and the development of consciousness (Hobson, 2009); however, the exact mechanisms and functions of REM sleep are far from unequivocal (Siegel, 2011).

In contrast to the prevailing view that conceptualizes REM sleep as a unified sleep stage, studies suggest that REM sleep is a heterogeneous state that is composed of (at least) two distinctive sub-states: phasic and tonic periods (De Carli et al., 2016; Simor et al., 2018). Phasic microstates with ocular movements, muscle twitches and vegetative irregularities are interspersed among longer and apparently more quiescent tonic periods. Several studies indicate that phasic and tonic REM sleep microstates are markedly different neural states as regarding awakening thresholds (Ermis et al., 2010), spontaneous oscillatory activity (Simor et al., 2016, 2018), information processing (Sallinen et al., 1996; Andrillon et al., 2017), and even mental experiences (Molinari and Foulkes, 1969; Stuart and Conduit, 2009).

Previous studies examining spontaneous oscillatory activity during phasic and tonic REM periods consistently found elevated levels in alpha (8 J 13 Hz) and beta (14 J 30 Hz) EEG power (Waterman et al., 1993; Jouny et al., 2000; De Carli et al., 2016; Simor et al., 2016) and synchronization (Simor et al., 2018) during tonic periods. These data suggest that tonic REM is closer to quiet wakefulness with regard to cortical oscillations. Spectral power and cortical synchronization in alpha-and beta frequency ranges measured during resting wakefulness is associated with a widespread cortical network that underpins the maintenance of alertness and facilitates the processing of external stimuli (Makeig and Inlow, 1993; Sadaghiani et al., 2010). Accordingly, reactivity to external (auditory) stimulation seems to be partially preserved during tonic states in contrast to phasic periods that appear as a functionally isolated state (Wehrle et al., 2007), detached from the environment (Sallinen et al., 1996; Ermis et al., 2010). On the other hand, relatively higher (> 31 Hz) gamma band power distinguished phasic from tonic microstates (Jouny et al., 2000; Abe et al., 2008; Simor et al., 2016). Enhanced gamma activity during phasic microstates was assumed to reflect sensorimotor, emotional and cognitive processes leading to intense dreaming (Gross and Gotman, 1999; Abe et al., 2008; Corsi-Cabrera et al., 2016).

Exploring the microarchitecture of REM sleep may provide novel insights into the mechanisms and functions of REM sleep; however, studies on REM microstates are still scarce, and more importantly, involve the data of usually few participants assessed with a low number of electrodes. Thus, the topographical aspects, neural sources, and patterns of functional connectivity of spontaneous cortical oscillations during REM microstates remain elusive, limiting our conclusions about the neurophysiological background of phasic and tonic REM sleep. Here, we analyzed high-density EEG (HD-EEG) recordings to contrast phasic and tonic REM periods with regard to the topographical distribution, source localization, as well as local, global and long-range synchronization of frequency-specific cortical activity. Our systematic analyses provide novel insights into the neurophysiological aspects of phasic and tonic microstates, sharpening the paradoxical nature of REM sleep.

## Materials and Methods

### Participants and procedure

Data were collected as part of a larger, research project (NKFI NK 104481) on cortical maturation and cognition (Gerván et al., 2017). Night-time EEG was recorded from twenty young adults for three consecutive nights at the sleep research laboratory of the Pázmány Péter Catholic University in Budapest. Participants arrived in the laboratory on Friday for their first recordings, therefore, the second and third nights were always Saturday and Sunday. This constraint allowed for a relative uniformity across subjects in terms of “sleep pressure” and daytime social demands. Participants arrived at the laboratory in the evening, and, after trained laboratory assistants applied the electrodes for electroencephalography (EEG), electromyography (EMG) and electrooculography (EOG), went to sleep according to their preferred bedtimes, scheduled between 10.00 PM and 11.30 PM. They slept until they awoke spontaneously. Only recordings from the second night were analyzed for this study, as these were free from adaptation effects (first night) and experimental manipulation (third night). After exclusion of two participants with a large number of noisy channels, eighteen subjects remained (eight females, mean age = 21.19, SD = 0.43) for data analysis. Based on the effect sizes found in an earlier study on REM sleep microstates (Simor et al., 2016), this number of participants is appropriate to detect effects of similar size at alpha = 0.05 with reasonable confidence (power between 0.65 and 0.99; pwr package, Champely, 2015). All participants were screened for and excluded if they had any current or previous sleep or mental disorder, or any neurological or medical condition. Fulfillment of inclusion and exclusion criteria were verified by standardized questionnaires as well as personal interviews. The consumption of alcohol and drugs (except contraceptives) was not allowed on the days of, and the day before the recordings. Additionally, participants were asked not to drink coffee and to refrain from napping on the afternoons of the days they were sleeping at the laboratory. Recruitment took place via social media networks and the social networks of Pázmány Péter Catholic University’s employees. All participants were right handed, and they received 20.000 HUF (approximately 70 US dollars) for their participation.

### Polysomnographic recording and preprocessing

Quick Cap 128 EEG caps (in three different sizes) with 128 built-in, passive Ag/AgCl electrodes (Compumedics, Australia) were used for recording EEG signals. These monopolar channels were referenced to a frontocentrally placed ground electrode. Bipolar EOG and EMG channels were also applied to measure eye movements and muscle tone, respectively. The EOG channels were placed below and to the left of the left eye (1 cm below the left outer canthus), and above and to the right of the right eye (1 cm above the right outer canthus). The EMG electrodes were applied to the chin symmetrically. The data was recorded using a BQ KIT SD LTM 128 EXPRESS (2 × 64 channels in master and slave mode, Micromed, Mogliano Veneto, Italy) recording device, and was subsequently visualized, saved and exported by the System Plus Evolution software (Micromed, Mogliano Veneto, Italy). All data (HD-EEG, EOG and EMG) was recorded with an effective sampling rate of 4096 Hz/channel (synchronous) with 22 bit equivalent resolution, prefiltered and amplified with 40 dB/decade high- and low-pass input filters (0.15-250 Hz). A digital low-pass filter was applied at 231.7 Hz, and the data was downsampled to 512 Hz/channel by firmware.

Sleep stages were scored manually according to standardized criteria (Berry et al., 2012) by experts trained in sleep research. From the epochs assigned to REM sleep, tonic and phasic REM sleep segments were selected manually by visual inspection in a custom-made software tool developed for all-night EEG data analysis (FercioEEGPlus, Ferenc Gombos, 2008-2018). Four second long segments were coded as phasic periods when within this time window the EOG channel showed at least two consecutive eye movements with 100 μV (or larger) amplitude. Four second long segments were coded tonic periods when no significant eye movements could be detected (EOG deflection of less than 25 μV) within the time window. To avoid contamination between the two microstates, segments were only selected if they were at least 8 seconds apart from each other and contained no movement-related or technical artifacts.

Preprocessing and further analysis of the data was done in MATLAB (version 9.3.0.713579, R2017b, The MathWorks, Inc., Natick, MA) using the open source toolbox Fieldtrip (Oostenveld et al., 2011). The code for all analyses can be found on OSF (Open Science Framework: *https://osf.io/tzvmk/?view_only=be7ce43fda0c48e7b01cf5d745fc68b2*). All data was re-referenced to the average of the mastoid channels, or, if either of these was noisy, to the average of TP7 and TP8 (in 7 of 18 cases). A band-pass filter of 0.5 to 70 Hz was applied, and line noise at 50 and 100 Hz was removed using a discrete Fourier transform filter. To increase the stationarity of the data, each segment was cut into 2 second trials with 50% overlap between them. As many more tonic than phasic segments were present in the data, the same number of available phasic segments was randomly selected from all tonic segments for each participant. Noisy channels were interpolated using the weighted average of surrounding electrodes, except when two or more neighboring channels were noisy, in which case an unweighted average of neighboring channels was used (in 9 of 18 cases). Trials were manually checked for artifacts, and contaminated trials were removed.

We applied independent component analysis (ICA) to identify and remove eye-movement related artifacts in EEG channels. Eye-movement artifacts have been found to contaminate the EEG signal measured from the scalp, especially in the lower frequency ranges (e.g. Croft and Barry, 2000). Moreover, others (Craddock et al., 2015) showed that saccade related eye muscle activity also confounds oscillatory activation in the gamma range. Performing ICA correction on the data has been found to successfully remove eye-movement related artefacts in both lower and higher frequency bands (Romero et al., 2003; Craddock et al., 2015). Tonic and phasic trials were appended for each participant, and EOG, EMG and reference channels were removed from the data before running the analysis. Components were identified as reflecting eye movement related artifacts if they showed a dominantly frontal topography and EOG-like time course. For most participants, two components showed this pattern and were therefore removed. In two participants, however, one or two additional components were identified and removed, while another participant had a component with large cardiac artifacts, which was also discarded.

A Laplacian transformation was performed on the data to improve the spatial precision of the spectral analysis and avoid spurious connectivity influences in measures of synchronization (Global Field Synchronization). By essentially acting as a spatial filter, Laplacian transformation of the data reduces the effect of volume conduction, as well as muscle and micro-saccade artifacts, and resolves reference issues, which are all known to influence EEG topography recorded from the scalp (Trujillo et al., 2005; Srinivasan et al., 2007; Tenke and Kayser, 2012) We followed the procedure introduced by Perrin and colleagues (Perrin et al., 1989) that calculates the Laplacian based on the 2^nd^ order spatial derivatives using spherical spline interpolation. This method has been found to be especially effective when using a large number of electrodes (Babiloni et al., 1995; Tandonnet et al., 2005). The parameters were set to the default settings for 128 electrodes in Fieldtrip (order of splines = 4, max. degree of Legendre polynomials = 20, *λ* = 10^−5^).

### Spectral power analysis

Spectral power was calculated for each 1 Hz frequency bin between 2 and 48 Hz. Tapering was applied to the two-second-long, artifact-free, ICA corrected and Laplace transformed phasic and tonic EEG segments for low and high frequencies separately: a Hanning taper for the lower frequencies (2-30 Hz) and a multi-taper using the dpss method with ± 2 Hz smoothing for the higher frequencies (31-48 Hz; Babapoor-Farrokhran et al., 2017). Power spectral density was computed for each participant, frequency bin and channel with the Fast Fourier Transformation (FFT) algorithm as implemented in Fieldtrip.

### Functional connectivity

Two separate measures of functional connectivity were used to examine differences in neural synchronization between phasic and tonic REM sleep periods. First, we calculated the Global Field Synchronization (GFS, Koenig et al., 2001), a measure reflecting the degree of phase alignment over all electrodes at a certain frequency. GFS was shown to efficiently characterize global EEG synchronization in different states of vigilance (Achermann et al., 2016) as well as to reveal impaired synchronization in pathological conditions (Koenig et al., 2001; Koenig et al., 2005). Segments of phasic and tonic REM sleep were transformed into the frequency domain using the same FFT algorithm and settings as for the spectral power analysis described above. This procedure decomposes the original signal into a real (the sine) and complex (the cosine) part (per each electrode and frequency). This way, all the EEG channels of each trial can be described as a cloud of points in a sine - cosine diagram. In order to quantify the amount of EEG activity that oscillate in a common phase in a given frequency, the procedure enters the sine and cosine values into a two-dimensional principal component analyses resulting in two eigenvalues (Koennig et al., 2005). GFS scores are computed as the ratios of these eigenvalues as defined by Koenig and colleagues (Koenig et al., 2001). We computed GFS for each frequency in each trial and subsequently averaged GFS values over trials for each participant. As can be understood from the formula, GFS values range from 0 to 1, with higher values indicating a higher degree of phase alignment over electrodes, thus implying more global synchronization.

In order to measure synchronization in local and large-scale neuronal ensembles, connectivity between short and long-range channel pairs was calculated using the Weighted Phase Lag Index (WPLI). WPU complements GFS as it is not sensitive to synchronized oscillations with zero phase lag (caused by a common generator or volume conduction), whereas GFS can be inflated by such oscillations. The Phase Lag Index was first introduced by Stam and colleagues (Stam et al., 2007). In essence, this method quantifies synchronization by the consistency of phase lags between two signals, using the distribution of phase angle differences. In contrast to other connectivity measures that are sensitive to volume conduction and common sources, the PLI attenuates the effect of volume conduction by disregarding phase angle differences at and around zero and *π*. The WPLI is a modified version of the PLI that takes into account that phase lags can turn into leads, and vice versa, by weighting phase angle differences around 0.5*π* and 1.5*π* more than those around zero and *π* (Vinck et al., 2011). This resolves the sensitivity of the PLI to small disturbances in phase lags, which is especially relevant for small synchronization effects. As WPLI is insensitive to volume conduction (Cohen, 2015), we used the raw scalp data (without Laplacian transformation) to calculate WPLI for each trial and 1 Hz frequency bin between 2 and 48 Hz, for all short and long range channel pairs. Short range channel pairs were within 5 cm from each other, while long range channel pairs were at least 20 cm apart. WPLI values fall between 0 and 1, with higher values indicating higher synchronization.

### Source reconstruction analysis

To explore the potential neural sources of the differences in spectral power across phasic and tonic microstates, beamformer source reconstruction analyses were performed for frequency bands that significantly differentiated the two conditions. The principal of beamformer techniques is to scan the brain location by location and estimate the proportion of the EEG signal measured at the scalp originating from each point using adaptive spatial filters (Gross et al., 2001). First, a model of the participants’ head is made, ideally based on individual anatomical scans. As no such scans were available for the participants in this study, the same standardized head model was used for all participants. This head model was based on a standard template MRI (Colin27) and constructed using a realistic boundary-element method (Oostenveld et al., 2003). Using this model and the positions of the sensors relative to the model of the head, a leadfield matrix was calculated for all points (sources) in the brain to be scanned. In the current study, the sources to be considered were defined by a regular 3D grid with a 5 mm resolution, thereby creating 3D voxels of 5 mm^3^. As beamforming techniques have been found to be ineffective for reliably reconstructing deep sources of EEG signals measured at the scalp (Muthuraman et al., 2014), only grid points located in the cortex were included in the source reconstruction analysis. Whether grid points were located in cortical areas or not was determined using the AAL atlas developed by (Tzourio-Mazoyer et al., 2002). Next, a common spatial filter was constructed for all grid points using the leadfield and the covariance matrix of the combined phasic and tonic data. This common filter was subsequently used to estimate the origin of the scalp-level spectral power for the tonic and phasic segments separately. The specific beamformer approach used in this study was Dynamic Imaging of Coherent Sources (DICS), as this method is suited for the localization of (differences in) spectral power (Gross et al., 2001). Since beamformer source reconstruction includes spatial filtering, preprocessed, ICA corrected data (without Laplacian transformation) was used to obtain spectral power as input for the beamformer analysis. We applied the same procedure for power analysis as described earlier, only on narrower frequency ranges. The regularization parameter was set at *λ* = 0.1% (Fuchs, 2007).

### Experimental Design and Statistical Analysis

All statistical analyses tested for differences between phasic and tonic REM sleep periods in a within-subject design.

Differences in spectral power and source localization were tested using cluster-based permutation tests suitable for the statistical analyses of HD-EEG data (Maris and Oostenveld, 2007). Two-sided paired samples t-tests were performed for all data points (i.e. frequency bins and electrodes for spectral power, and source locations for reconstructed sources), and clustered if adjacent frequencies/locations showed significant differences at ⍰ < 0.05. For the analysis on power differences, at least four neighboring electrodes had to be significant to be considered as a cluster. We used this additional criterion due to the large number of electrodes used in this study. A cluster statistic was calculated for each identified cluster by summing the t-values belonging to the included data points. The same process was repeated 1000 times, but with phasic and tonic trials randomly shuffled within participants using the Monte-Carlo method implemented in Fieldtrip (Maris and Oostenveld, 2007). The largest cluster statistic of each permutation was used to create the probability distribution against which the observed cluster statistics were tested with an alpha value of 0.05. Cluster-level corrected p-values are reported throughout the text. No commonly accepted measure of effect size for cluster-based permutation tests on high dimensional EEG data has been established up to this point. Still, to give some indication of effect size, Cohen’s d was calculated on the averaged data within clusters. This measure of effect size should be considered with caution, however.

For both synchronization analyses, bootstrap statistics were used to test for differences between phasic and tonic REM sleep (Tarokh et al., 2011). WPLI values were averaged over short and long range channel pairs, and only differences between phasic and tonic periods were investigated (i.e. long and short range connectivity were not contrasted directly). For each frequency bin, a paired-samples t-test probability distribution was calculated based on samples taken from the original data, randomly shuffling phasic and tonic REM data within subjects (package mosaic in R; (Pruim et al., 2017). 10 000 samples were used to create the probability distributions, as this was observed to lead to an acceptably small amount of variation in p-values (± 0.009). The difference between phasic and tonic data was subsequently tested against this probability distribution with a two-tailed dependent samples t-test with an alpha value of 0.05. Following the example of Tarokh and colleagues (Tarokh et al., 2010), the findings were corrected for multiple comparisons by only reporting differences when at least three adjacent frequency bins showed significance, as the likelihood of this representing random activity is small. Effect sizes (Cohen’s d) were calculated with the bootES package in R (Kirby and Gerlanc, 2017).

## Results

### Sleep architecture

Conventional parameters of sleep architecture are presented in Table 1. Participants showed good sleep efficiency (Mean = 90.64 Std. Dev = 7.33) and on average spent 17.15 % and 26.41 % in Slow Wave Sleep and REM sleep, respectively. The sleep architecture of our sample closely corresponded to the norms of this age group (Carskadon and Dement, 2005).

**Table 1.** Conventional parameters of sleep architecture of the study sample.

|  | **Mean** | **Standard Deviation** |
| --- | --- | --- |
| Sleep efficiency (%) | 90.64 | 7.33 |
| Wake after sleep onset (min) | 18.58 | 31.83 |
| Sleep onset latency (min) | 32.83 | 25.7 |
| NREM duration (%) | 73.59 | 4.14 |
| Stage 1 duration (%) | 1.49 | 0.77 |
| Stage 2 duration (%) | 54.93 | 5.65 |
| Slow Wave Sleep duration (%) | 17.17 | 5.64 |
| REM duration (%) | 26.41 | 4.14 |
| REM latency (min) | 98.97 | 42.59 |

### Spectral power

The cluster-based permutation test revealed that phasic and tonic REM periods differed significantly from each other in spectral power. This difference was expressed in two large clusters: one negative cluster (indicating higher power in the tonic microstate) in the high alpha/beta bands (12-28 Hz, t_maxsum_ = −3738.9, p = .002, d = 1.97) and one positive cluster (indicating higher power in the phasic microstate) in the gamma frequency range (33-48 Hz, t_maxsum_ = 3490.8, p = .002, d = 2.05). An additional positive cluster (indicating higher power in the phasic microstate) in the delta band showed a trend (2-4 Hz, t_maxsum_ = 412.23, p = .056, d = 2.1). The topographic distribution of these clusters is shown in Figure 1. The negative cluster in the high alpha/beta frequency ranges was spread over frontocentral electrodes (marked by white asterisks), while the positive cluster in the gamma range extended to posterior as well as anterior and central regions (marked by black asterisks). The cluster in the delta range was mostly present in frontocentral sites (marked by black plus signs).

**Figure 1.**
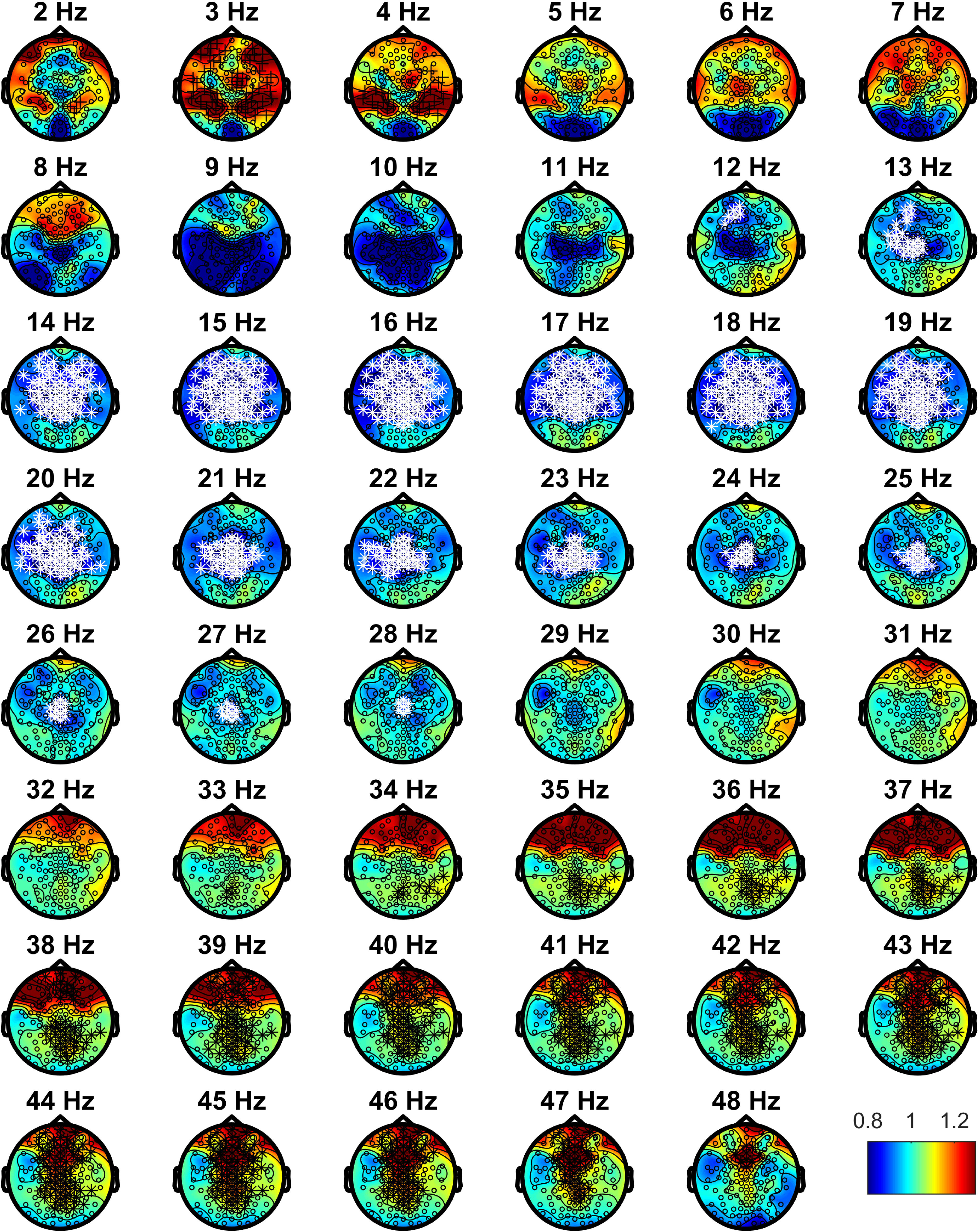
Contrast of absolute spectral power of phasic / tonic REM microstates in each examined frequency bin. Hot colors indicate relatively higher power in the phasic, and cold colors indicate relatively higher power in the tonic microstate. Black plus signs mark the positive cluster showing a trend in the delta band, white asterisks mark the negative cluster in the high alpha/beta band, black asterisks mark the positive cluster in the gamma band. Positive clusters indicate relatively higher spectral power during phasic REM sleep, whereas negative clusters indicate relatively higher spectral power during tonic REM sleep.

### Global Field Synchronization

Bootstrap statistics run on the GFS values showed significant differences in global synchronization between phasic and tonic microstates in the delta, theta and beta range (all p < .021, d = 0.52-1.30). In the delta and theta range, global synchronization was increased in phasic compared to tonic REM, while the opposite was true in the beta range. The significant frequency bins are indicated by asterisks in Figure 2 (if at least three adjacent frequency bins were significant).

**Figure 2.**
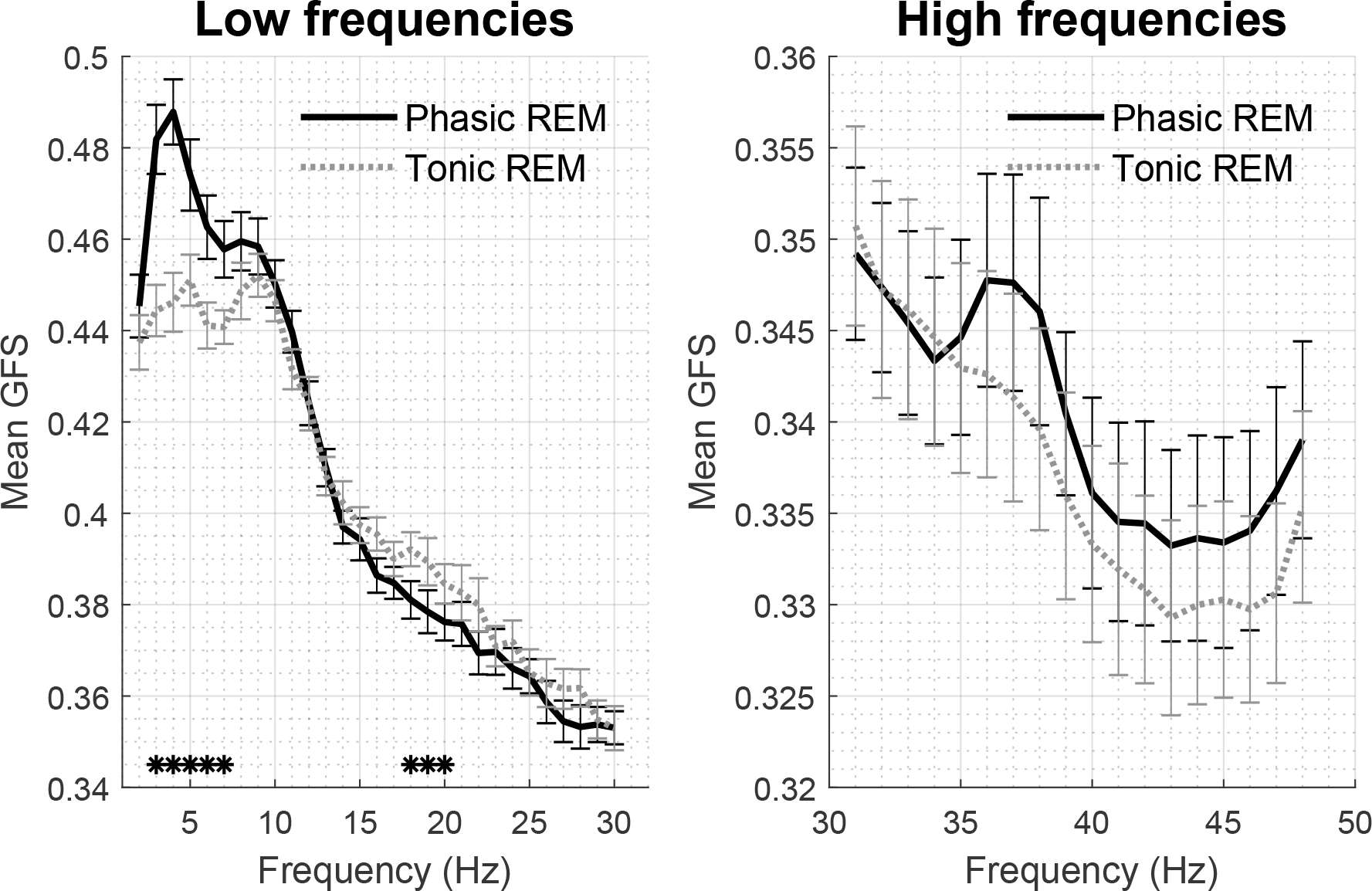
Mean Global Field Synchronization during tonic and phasic REM sleep for low (left) and high (right) frequencies. Asterisks mark significant differences as identified by the bootstrap procedure.

### Long and short range synchronization

Patterns of long and short range WPLI values in phasic and tonic states are illustrated in Figure 3. For both long and short range WPLI, bootstrap statistics revealed significant differences between phasic and tonic REM periods at several frequencies. More specifically, synchronization was elevated for at least three consecutive frequency bins in phasic compared to tonic REM sleep at the delta (2-4 Hz for both long and short range, all p < .011, d = 0.58-1.26) and gamma ranges (31-48 Hz in case of short range WPLI, all p < .001; 37-47 Hz in case of long range WPLI, all p < .017, d = 0.5-2.3). WPLI in single frequency bins corresponding to the alpha and beta ranges showed trends of increased synchrony in tonic periods in both short and long distance pairs; however, these differences did not remain significant after the correction for multiple comparisons. WPLI values as well statistically significant differences across phasic and tonic states are presented in Figure 3.

**Figure 3.**
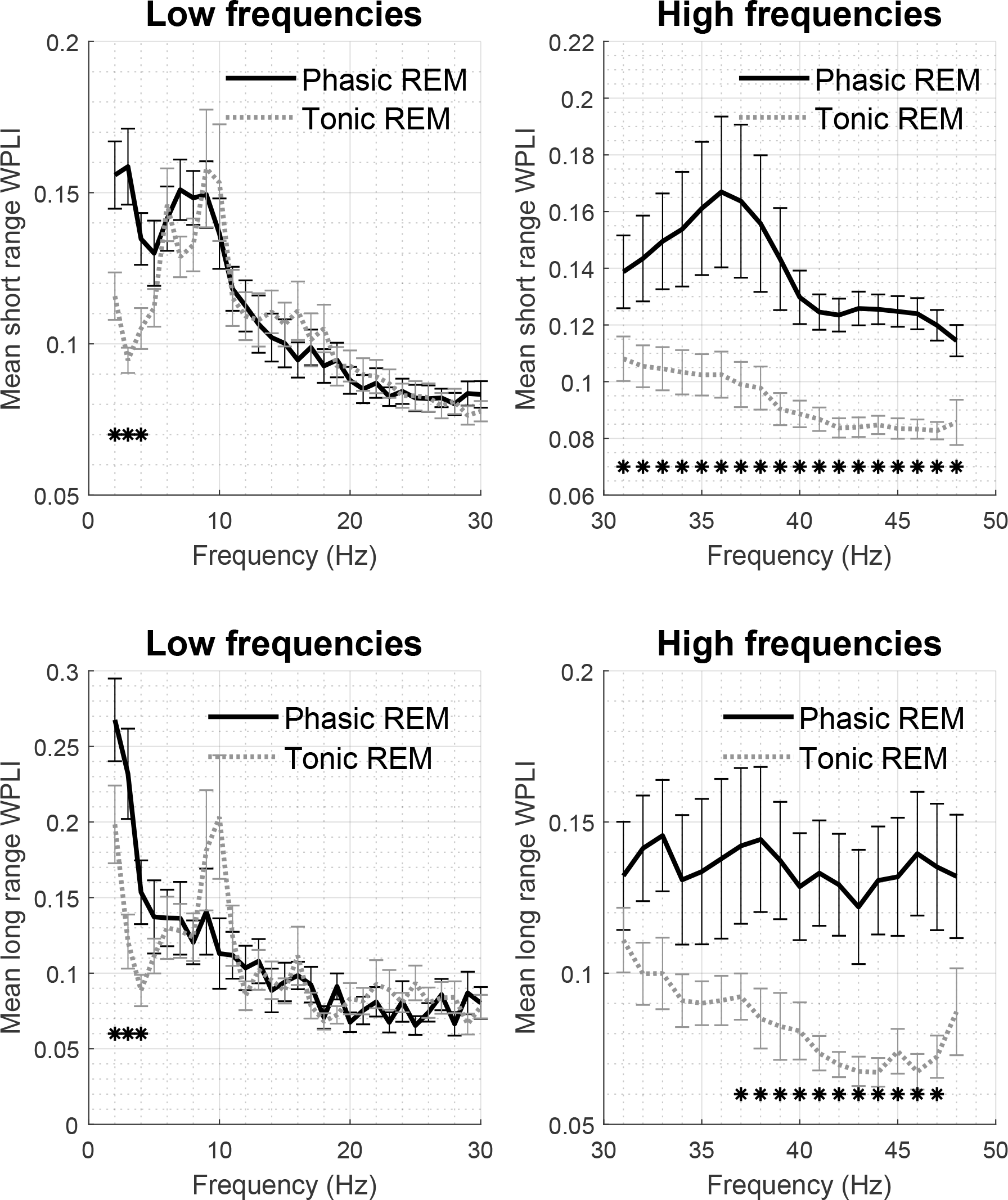
Mean short range (on the top) and long range (on the bottom) WPLI during phasic and tonic REM sleep for low (left) and high (right) frequencies. Asterisks mark significant differences as identified by the bootstrap procedure.

### Source reconstruction

Beamformer source reconstruction analyses were performed in the frequency ranges that distinguished phasic from tonic states in terms of spectral power (high alpha/beta frequencies between 12-28 Hz and gamma frequencies between 33-48 Hz). In case of the high alpha/beta ranges, the cluster-based permutation test of the identified sources revealed a significant difference across phasic and tonic microstates expressed in one negative cluster (t_maxsum_ = −9754, p = .002, d = 0.88, cluster-level corrected). The negative direction of this cluster indicates that spectral power was higher during tonic compared to phasic REM sleep in the identified sources. This cluster was widely spread over the central areas of the brain, including bilateral areas around the central sulcus (including motor and somatosensory areas) and around the temporo-parietal junction (e.g. supramarginal and angular gyri). The areas found to belong to the cluster are listed in Table 2. Two positive clusters were identified as potential sources of significantly higher gamma power in phasic as compared to tonic REM (t_maxsum_ = 2786, p = .01, d = 0.98; t_maxsum_ = 1391.4, p = .03, d = 0.80, cluster-level corrected). The areas belonging to the first cluster were located mainly in bilateral medial frontal regions, such as the medial frontal gyrus and the orbital part of the middle frontal gyrus. Areas of the second cluster were primarily located in the right temporal lobe, including regions like the fusiform gyrus and superior, middle and inferior temporal gyrus, but also included some areas in the parietal (e.g. posterior cingulate gyrus and precuneus) and occipital (e.g. lingual gyrus and calcarine sulcus) lobes. The full lists of areas belonging to these clusters are presented in Table 3 and Table 4. The tables are sorted according to the percentage of voxels forming part of the identified clusters, compared to the total number of voxels included in the analysis for each area, as determined with the AAL atlas. The t-values of the voxels belonging to all three clusters are plotted onto a normalized brain surface based on a standardized MRI model in Figure 4.

**Table 2.** AAL labels of the areas containing voxels that belong to the negative cluster indicating significantly higher power in the high alpha/beta range during tonic compared to phasic REM sleep. The percentage of voxels is calculated based on the total number of voxels present in that area.

| **AAL label** | **Number of voxels** | **Percentage of voxels** |
| --- | --- | --- |
| Heschl's gyrus (L) | 16 | 100.00% |
| Supramarginal gyrus (L) | 67 | 98.53% |
| Rolandic Operculum (L) | 50 | 98.04% |
| Inferior Parietal lobule (L) | 147 | 98.00% |
| Postcentral gyrus (L) | 251 | 97.67% |
| Precentral gyrus (L) | 211 | 94.20% |
| Inferior Frontal gyrus, pars opercularis (L) | 57 | 93.44% |
| Superior Parietal lobule (R) | 119 | 92.97% |
| Superior Parietal lobule (L) | 110 | 88.71% |
| Superior Temporal gyrus (L) | 120 | 88.24% |
| Paracentral lobule (L) | 74 | 88.10% |
| Paracentral lobule (R) | 48 | 87.27% |
| Postcentral gyrus (R) | 167 | 71.37% |
| Precentral gyrus (R) | 146 | 70.19% |
| Supplementary Motor Area (L) | 93 | 66.91% |
| Insula (L) | 66 | 65.35% |
| Angular gyrus (L) | 43 | 62.32% |
| Frontal Inf Tri (L) | 95 | 60.90% |
| Inferior Parietal lobule (R) | 53 | 59.55% |
| Superior Frontal gyrus (L) | 104 | 48.37% |
| Precuneus (R) | 93 | 47.94% |
| Precuneus (L) | 100 | 47.85% |
| Superior Frontal gyrus (R) | 115 | 46.94% |
| Supplementary Motor Area (R) | 68 | 46.90% |
| Middle Frontal gyrus (L) | 142 | 46.71% |
| Midcingulate area (L) | 40 | 34.48% |
| Middle Frontal gyrus (R) | 102 | 33.89% |
| No label found | 110 | 32.64% |
| Middle Temporal gyrus (L) | 98 | 30.34% |
| Midcingulate area(R) | 30 | 24.79% |
| Supramarginal gyrus (R) | 19 | 17.12% |
| Superior Temporal gyrus (R) | 32 | 16.58% |
| Inferior Frontal gyrus, pars opercularis (R) | 12 | 14.29% |
| Medial Frontal gyrus (L) | 27 | 13.24% |
| Superior Temporal Pole (L) | 10 | 12.50% |
| Medial Frontal gyrus (R) | 17 | 11.49% |
| Superior Occipital gyrus (R) | 8 | 9.88% |
| Angular gyrus (R) | 10 | 9.71% |
| Cuneus (R) | 5 | 5.32% |
| Anterior cingulate gyrus (L) | 5 | 4.85% |
| Inferior Frontal gyrus, pars orbitalis (L) | 3 | 2.78% |
| Middle Occipital gyrus (L) | 5 | 2.46% |
| Middle Temporal gyrus (R) | 5 | 1.82% |
| Rolandic Operculum (R) | 1 | 1.19% |

**Table 3.**
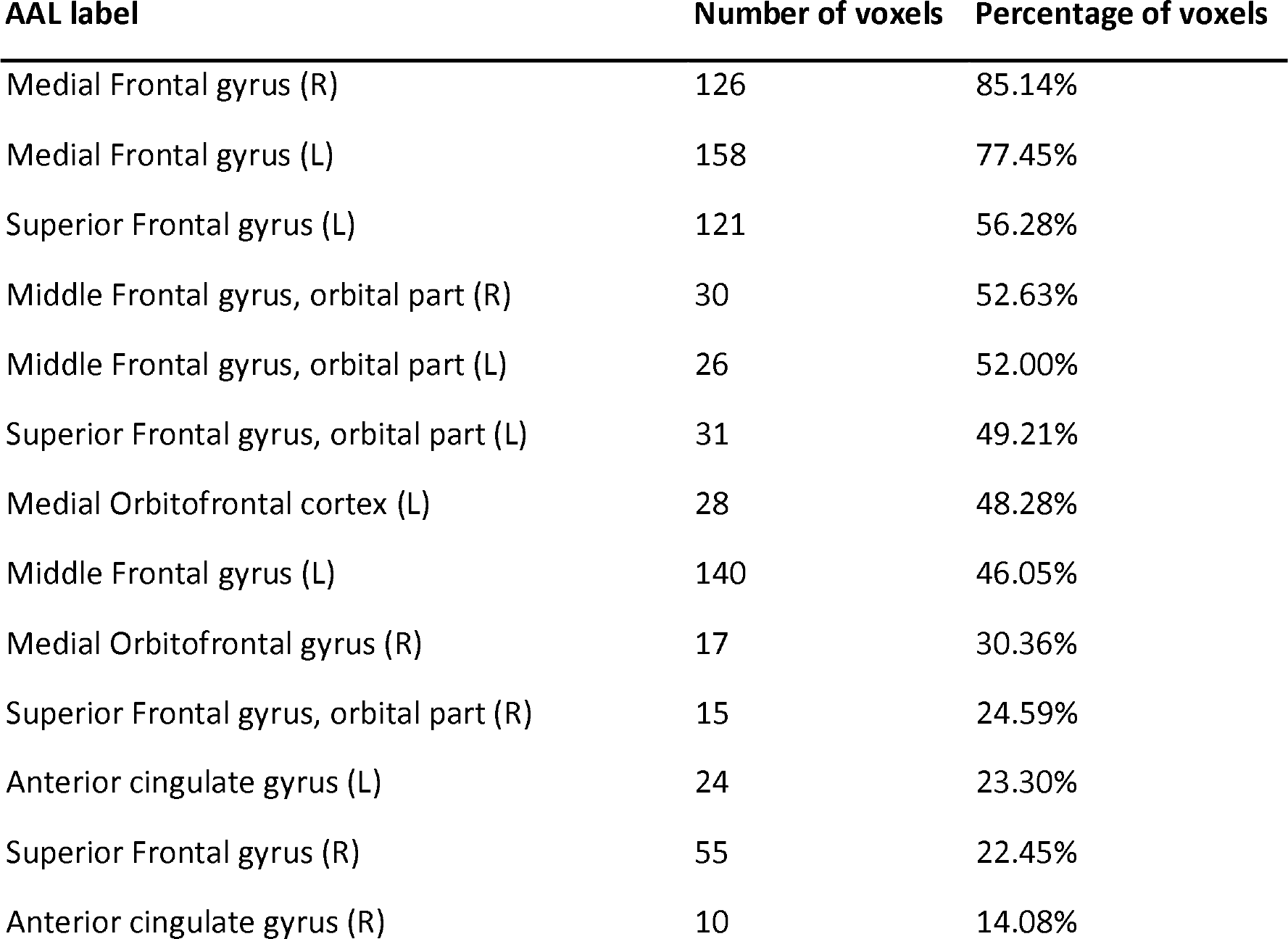
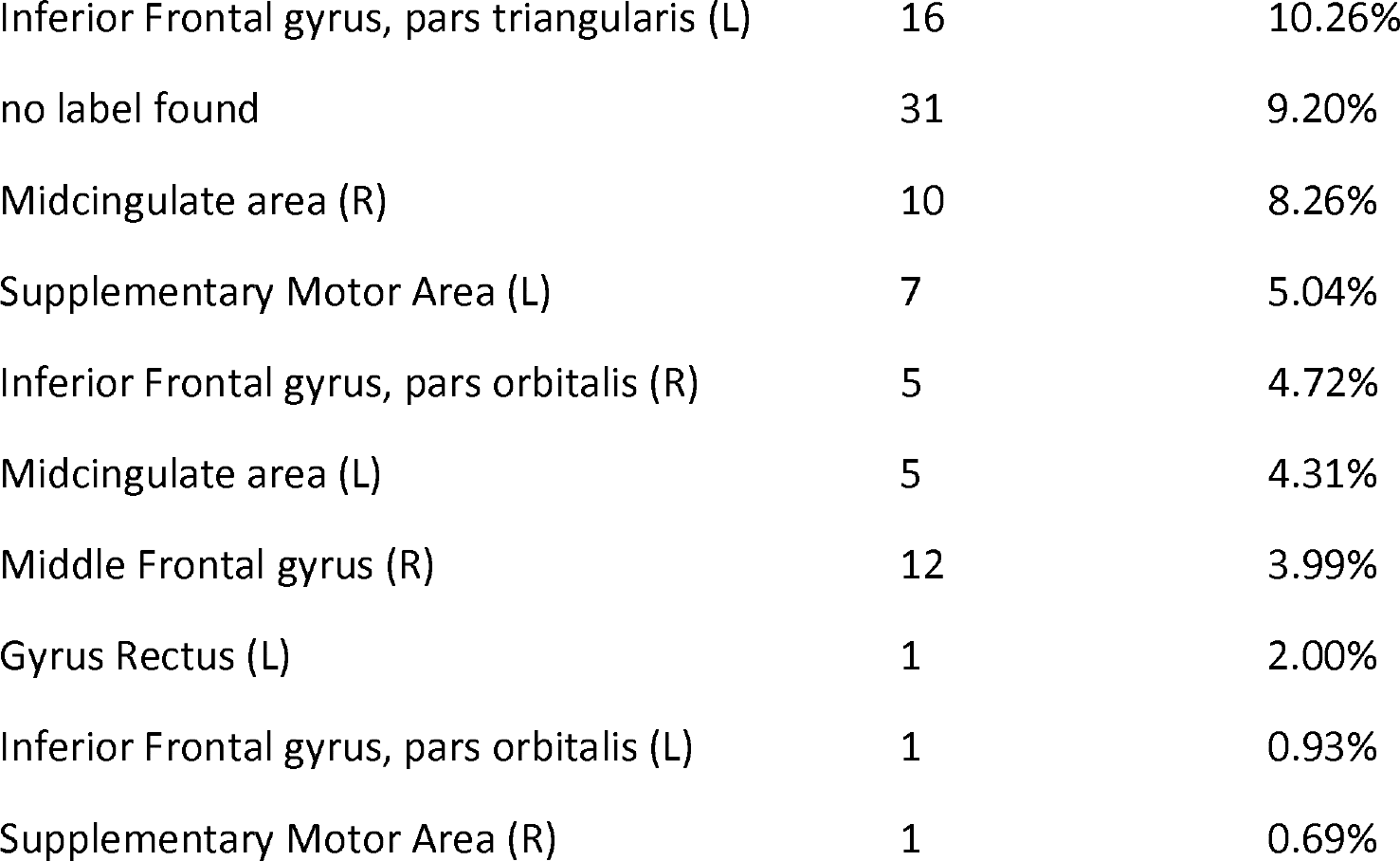
AAL labels of the areas containing voxels that belong to the first positive cluster indicating significantly higher power in the gamma frequency range during phasic compared to tonic REM sleep. The percentage of voxels is calculated based on the total number of voxels present in that area.

**Table 4.** AAL labels of the areas containing voxels that belong to the second positive cluster indicating significantly higher power in the gamma frequency range during phasic compared to tonic REM sleep. The percentage of voxels is calculated based on the total number of voxels present in that area.

| **AAL label** | **Number of voxels** | **Percentage of voxels** |
| --- | --- | --- |
| Fusiform gyrus (R) | 101 | 65.58% |
| Heschl's gyrus (R) | 9 | 64.29% |
| Posterior Cingulate gyrus (R) | 11 | 44.00% |
| Superior Temporal gyrus (R) | 82 | 42.49% |
| Middle Temporal gyrus (R) | 89 | 32.48% |
| Inferior Temporal gyrus (R) | 52 | 24.76% |
| Lingual gyrus (R) | 32 | 23.36% |
| Calcarine sulcus (R) | 24 | 22.02% |
| Precuneus (R) | 29 | 14.95% |
| Insula (R) | 16 | 14.04% |
| Rolandic Operculum (R) | 9 | 10.71% |
| No label found | 24 | 7.12% |
| Middle Temporal Pole (R) | 3 | 4.76% |
| Angular gyrus (R) | 2 | 1.94% |
| Cuneus (R) | 1 | 1.06% |

**Figure 4.**
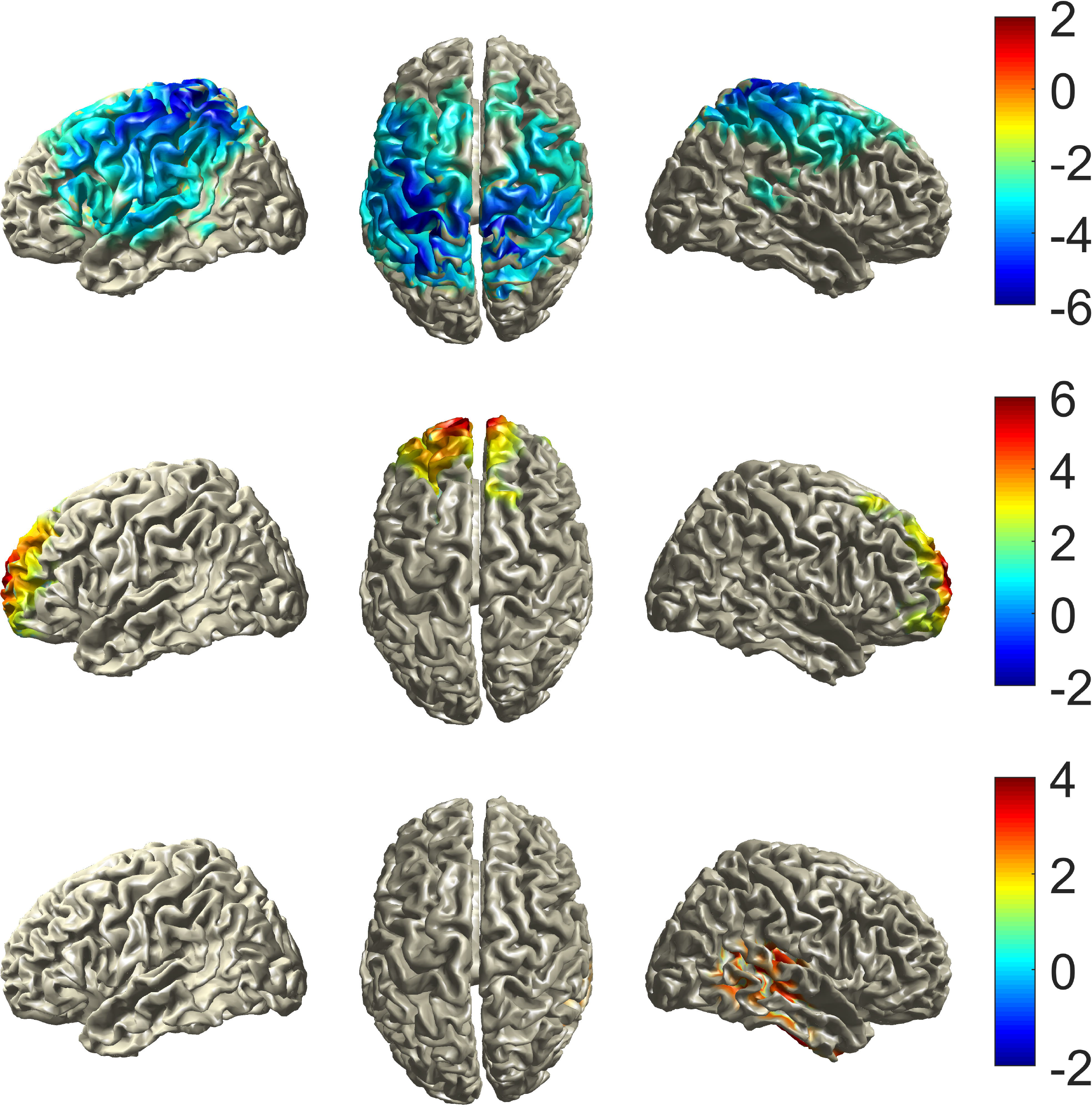
Statistical maps (significant t-values) plotted on top of a standardized brain surface for the high alpha/beta cluster (top), the first gamma cluster (middle) and the second gamma cluster (bottom).

## Discussion

Our aim was to examine phasic and tonic REM microstates in terms of cortical oscillations, topography, and neural synchronization by analyzing full night HD-EEG recordings of young healthy individuals. We showed that high alpha and beta power is relatively increased during tonic REM sleep and widely expressed over frontocentral locations. On the other hand, high frequency gamma power along the anterior-posterior axis, and to a lesser extent, low frequency oscillations were more prominent during phasic periods. Source localization of high alpha/beta and gamma band power revealed several cortical sites as potential generators that contributed differently to express such oscillations in phasic and tonic REM sleep. Beyond differences in spectral power, low frequency (3-7 Hz) oscillations in the delta and theta ranges showed higher global synchronization during phasic, whereas oscillations in the beta range exhibited enhanced global synchronization during tonic periods. Furthermore, low (2-4 Hz) frequency and gamma oscillations showed increased WPLI during phasic compared to tonic states in case of both short and long distance electrode pairs.

Our findings corroborate and extend previous studies that reported increased low frequency power during phasic compared to tonic REM sleep (Waterman et al., 1993; Jouny et al., 2000; Simor et al., 2016). One plausible interpretation is that delta power reflects eye movements that contaminate EEG signals. While previous studies could not appropriately minimize the effects of eye movement related artifacts, in the present study we may exclude this possibility. First, we attenuated eye movement artifacts by ICA. Secondly, we applied Laplacian transformation to minimize further the influence of electric potentials generated by the eyes. We only considered oscillations above 2 Hz, as eye movement potentials mainly extend over 0.3 and 2 Hz (Tan et al., 2001). Moreover, eye movement artifacts are usually restricted to frontal derivations (Waterman et al., 1992), but in our study increased slow frequency power in phasic REM was spread (at a trend level) over more posterior regions, arguing against a mere effect of ocular contamination. Finally, slow frequency activity showed increased synchronization as measured by GFS and WPLI. Although the former is sensitive to volume conduction, the latter discards phase couplings of zero (or 180°) phase lag, and hence, is immune to ocular artifacts transmitted by volume conduction.

Low frequency (1-4 Hz) oscillations are the hallmark of Slow Wave Sleep (SWS) and emerge from thalamo-cortical and cortico-cortical networks, in which widely distributed neurons rhythmically alternate between hyperpolarized and depolarized states (Steriade et al., 1993). Low frequency oscillations seem to underlie sensory disconnection and promote the restorative functions of sleep (Borbély, 1982; Tononi and Cirelli, 2006). Nevertheless, low frequency oscillations are not limited to SWS, but can also appear in wakefulness (Vyazovskiy et al., 2011; Fattinger et al., 2017) and REM sleep (Funk et al., 2016; Baird et al., 2018), especially in case of high homeostatic sleep pressure (Marzano et al., 2010). As awakening thresholds (Ermis et al., 2010) and auditory processing (Wehrle et al., 2007) are reduced in phasic conditions, we suggest that relatively increased and widely synchronized low frequency oscillations in phasic compared to tonic periods reflect a deeper sleep state, that is disconnected from the environment.

Relatively increased and more synchronized low frequency oscillations during the phasic as compared to the tonic state suggest that the former is closer to deep sleep; however, low frequency activity in phasic REM appears to coincide with increased high frequency oscillations reflecting intense cortical activity and resembling wakefulness (Cantero et al., 2004). Increased gamma power in phasic periods was consistently found in previous scalp EEG (Jouny et al., 2000; Simor et al., 2016) and intracerebral measurements (Gross and Gotman, 1999). Here we showed that phasic gamma activity is remarkably widespread over the scalp and exhibits higher synchronization between short and long distance electrode locations. Gamma oscillations in phasic periods showed significantly stronger synchronization by the WPLI, but were not different across microstates in terms of GFS. This discrepancy indicates that gamma synchrony does not emerge from a common generator or widespread cortical ensembles, but may reflect the activity of more localized networks.

Source localization unfolded two clusters as potential sources of increased gamma power in phasic periods. The first consisted of regions that correspond to the superior and ventromedial prefrontal cortex (e.g. medial frontal and orbitofrontal gyrus, anterior cingulate). This findings is in line with neuroimaging studies reporting the selective activation of frontolimbic networks following REMs (Muzur et al., 2002; Wehrle et al., 2007; Miyauchi et al., 2009). Phasic activity such as pontine waves in rodents (Datta and O’Malley, 2013), as well as frontolimbic activity in relation to REM sleep (Spoormaker et al., 2014) were linked to emotional regulation (more specifically, fear extinction), a process often associated with REM sleep in general (Levin and Nielsen, 2007; Genzel et al., 2015). The second cluster of sources comprised right lateralized temporal and posterior regions including the superior temporal, lingual, Heschl’s and fusiform gyrus. These regions form important pathways between the amygdala and prefrontal structures, and play a key role in perceptual processes, especially in the perception of emotional stimuli (Radua et al., 2010; Kawasaki et al., 2012). In a recent study, Corsi-Cabrera and colleagues (2016) showed that the human amygdala is generating gamma oscillations immediately after REMs. Therefore, we may speculate that increased and widely synchronized gamma oscillations during phasic REM sleep might (partly) originate from activity of the amygdala and related frontolimbic structures underlying emotional, perceptual and regulatory (e.g. fear extinction) processes that humans experience in the form of dream images.

Previous studies consistently reported relatively increased (high) alpha and beta power during tonic compared to phasic periods (Jouny et al., 2000; De Carli et al., 2016; Simor et al., 2016). Our findings with HD-EEG extend these data by showing that increased high alpha and beta power in tonic REM are most prominent at frontocentral derivations, but exhibit a widespread pattern over the scalp between 14 and 20 Hz. Interestingly, frequency bins within the beta band showed relatively increased GFS during tonic periods, but WPLI did not conclusively reveal higher synchronization of beta frequencies in tonic compared to phasic periods. (Of note, WPLI showed trends in high alpha and beta frequencies in the expected direction, but these differences were not significant after correction for multiple comparisons). In a recent study, Simor and colleagues (2018) suggested that enhanced high alpha and beta activity reflect increased environmental alertness during tonic REM sleep. External processing appears to be partially maintained during tonic states in contrast to phasic periods (Sallinen et al., 1996; Wehrle et al., 2007; Andrillon et al., 2017), and high alpha and beta power in resting wakefulness is associated with a neural network that maintains alertness, and facilitates the processing of external stimuli (Sadaghiani et al., 2010). Nevertheless, it is still premature to infer that high alpha and beta oscillations during tonic REM directly reflect such cognitive processes.

As regarding the sources of increased high alpha/beta power during tonic microstates, we identified several regions corresponding to higher-order, multimodal, associative areas, such as the supramarginal gyrus, the inferior parietal and inferior frontal lobules, as well as the superior frontal gyrus (Jung et al., 2017). In addition, a potential source in the left Heschl’s gyrus emerged, a region that is crucial in auditory processing (Warrier et al., 2009). Sensorimotor areas also emerged as sources of high alpha/beta power in line with an intracranial study showing relatively increased alpha/beta power during tonic states in the primary motor cortex resembling the activity found in relaxed wakefulness. In contrast, reduced alpha/beta power in the motor cortex during phasic REM was similar to the activity found during executed movements (De Carli et al., 2016). Tonic REM sleep might thus resemble resting wakefulness, with higher-order cognitive processes, increased environmental awareness, and attenuated primary sensorimotor cortical activity.

In light of these data, it is tempting to speculate that during tonic states the brain favors higher-order thought processes, whereas during phasic periods emotional and sensorimotor processes predominate. These may underlie the more active, perceptually vivid, and less thought-like dreams reports obtained after awakenings from phasic than after tonic REM periods (Molinari and Foulkes, 1969; Arnulf, 2011).

REM sleep was termed as a paradoxical state due to the combination of an activated cortex and a paralyzed body (Jouvet, 1965). Our study indicates that the paradox is even more complex. Phasic REM periods are characterized by the combination of synchronized low frequency oscillations resembling deeper sleep stages and high frequency oscillations resembling wake-like cortical activity. In contrast, tonic REM sleep is apparently a more quiescent state; however, increased oscillatory activity in the alpha and beta ranges and increased environmental alertness resembles neural processes of resting wakefulness.

Our study is the first to examine the neurophysiological aspects of phasic and tonic microstates with HD-EEG; however, future studies may improve spatial resolution by applying individual MR head models, or HD-EEG combined with Magnetoencephalography. Furthermore, studies in larger and more heterogeneous samples may reveal the influence of age and sex on neural activity during phasic and tonic REM microstates.

## Acknowledgements

The project was supported by the UNKP-18-4 (Bolyai +) New National Excellence Program of the Ministry of Human Capacities. P.S. was supported by the Hungarian Scientific Research Fund (NKFI FK 128100) of the National Research, Development and Innovation Office and by the Bolyai János Research Scholarship of the Hungarian Academy of Sciences. I. K. was supported by the Hungarian Scientific Research Fund (NKFI NK 104481) of the National Research, Development and Innovation Office.

## Notes

Conflict of Interest: The authors declare no competing financial interests.

## References

Abe T, Matsuoka T, Ogawa K, Nittono H, Hori T (2008) Gamma band EEG activity is enhanced after the occurrence of rapid eye movement during human REM sleep. Sleep Biol Rhythms 6:26–33.

Achermann P, Rusterholz T, Dürr R, König T, Tarokh L (2016) Global field synchronization reveals rapid eye movement sleep as most synchronized brain state in the human EEG. R Soc Open Sci 3:160201.

Andrillon T, Pressnitzer D, Léger D, Kouider S (2017) Formation and suppression of acoustic memories during human sleep. Nature Communications 8:179.

Arnulf I (2011) The “scanning hypothesis” of rapid eye movements during REM sleep: a review of the evidence. Arch Ital Biol 149:367–382.

Babapoor-Farrokhran S, Vinck M, Womelsdorf T, Everling S (2017) Theta and beta synchrony coordinate frontal eye fields and anterior cingulate cortex during sensorimotor mapping. Nature Communications 8:13967.

Babiloni F, Babiloni C, Fattorini L, Carducci F, Onorati P, Urbano A (1995) Performances of surface Laplacian estimators: A study of simulated and real scalp potential distributions. Brain Topogr 8:35–45.

Baird B, Castelnovo A, Riedner BA, Lutz A, Ferrarelli F, Boly M, Davidson RJ, Tononi G (2018) Human Rapid Eye Movement Sleep Shows Local Increases in Low-Frequency Oscillations and Global Decreases in High-Frequency Oscillations Compared to Resting Wakefulness. eNeuro:ENEURO.0293–18.2018.

Berry RB, Brooks R, Gamaldo CE, Harding SM, Marcus CL, Vaughn BV (2012) The AASM manual for the scoring of sleep and associated events. Rules, Terminology and Technical Specifications, Darien, Illinois, American Academy of Sleep Medicine Available at: http://www.aasmnet.org/resources/pdf/scoring-manual-preface.pdf [Accessed May 15, 2017],

Blumberg MS, Coleman CM, Gerth AI, McMurray B (2013) Spatiotemporal structure of REM sleep twitching reveals developmental origins of motor synergies. Curr Biol 23:2100–2109.

Borbély AA (1982) A two process model of sleep regulation. Hum neurobiol 1:195–204.

Cantero JL, Atienza M, Madsen JR, Stickgold R (2004) Gamma EEG dynamics in neocortex and hippocampus during human wakefulness and sleep. Neuroimage 22:1271–1280.

Carskadon MA, Dement WC (2005) Normal human sleep: an overview. Principles and practice of sleep medicine 4:13–23.

Champely S (2015) pwr: Basic functions for power analysis. R package version 1.

Cohen MX (2015) Effects of time lag and frequency matching on phase-based connectivity. J Neurosci Methods 250:137–146.

Corsi-Cabrera M, Velasco F, Del Río-Portilla Y, Armony JL, Trejo-Martínez D, Guevara MA, Velasco AL (2016) Human amygdala activation during rapid eye movements of rapid eye movement sleep: an intracranial study. J Sleep Res 25:576–582.

Craddock M, Martinovic J, Müller MM (2015) Accounting for microsaccadic artifacts in the EEG using independent component analysis and beamforming. Psychophysiology 53:553–565.

Croft RJ, Barry RJ (2000) Removal of ocular artifact from the EEG: a review. Neurophysiologie Clinique/Clinical Neurophysiology 30:5–19.

Datta S, O’Malley MW (2013) Fear extinction memory consolidation requires potentiation of pontine-wave activity during REM sleep. J Neurosci 33:4561–4569.

De Carli F, Proserpio P, Morrone E, Sartori I, Ferrara M, Gibbs SA, De Gennaro L, Lo Russo G, Nobili L (2016) Activation of the motor cortex during phasic rapid eye movement sleep. Ann Neurol 79:326–330.

Ermis U, Krakow K, Voss U (2010) Arousal thresholds during human tonic and phasic REM sleep. Journal of sleep research 19:400–406.

Fattinger S, Kurth S, Ringli M, Jenni OG, Huber R (2017) Theta waves in children’s waking electroencephalogram resemble local aspects of sleep during wakefulness. Sci Rep 7:11187.

Fuchs A (2007) Beamforming and its applications to brain connectivity. In: Handbook of Brain Connectivity, pp 357–378. Springer.

Funk CM, Honjoh S, Rodriguez AV, Cirelli C, Tononi G (2016) Local Slow Waves in Superficial Layers of Primary Cortical Areas during REM Sleep. Curr Biol 26:396–403.

Genzel L, Spoormaker VI, Konrad BN, Dresler M (2015) The role of rapid eye movement sleep for amygdala-related memory processing. Neurobiol Learn Mem 122:110–121.

Gervén P, Soltész P, Filep O, Berencsi A, Kovács I (2017) Posterior-Anterior Brain Maturation Reflected in Perceptual, Motor and Cognitive Performance. Front Psychol 8:674.

Gross DW, Gotman J (1999) Correlation of high-frequency oscillations with the sleep-wake cycle and cognitive activity in humans. Neuroscience 94:1005–1018.

Gross J, Kujala J, Hämäläinen M, Timmermann L, Schnitzler A, Salmelin R (2001) Dynamic imaging of coherent sources: Studying neural interactions in the human brain. PNAS 98:694–699.

Hobson JA (2009) REM sleep and dreaming: towards a theory of protoconsciousness. Nature Reviews Neuroscience 10:803–813.

Jouny C, Chapotot F, Merica H (2000) EEG spectral activity during paradoxical sleep: further evidence for cognitive processing. NeuroReport 11:3667–3671.

Jouvet M (1965) Paradoxical Sleep — A Study of its Nature and Mechanisms. Progress in Brain Research 18:20–62.

Jung J, Cloutman LL, Binney RJ, Lambon Ralph MA (2017) The structural connectivity of higher order association cortices reflects human functional brain networks. Cortex 97:221–239.

Kawasaki H, Tsuchiya N, Kovach CK, Nourski KV, Oya H, Howard MA, Adolphs R (2012) Processing of facial emotion in the human fusiform gyrus. J Cogn Neurosci 24:1358–1370.

Kirby KN, Gerlanc D (2017) Finding Bootstrap Confidence Intervals for Effect Sizes With BootES. APS Observer 30 (3).

Koenig T, Lehmann D, Saito N, Kuginuki T, Kinoshita T, Koukkou M (2001) Decreased functional connectivity of EEG theta-frequency activity in first-episode, neuroleptic-na’ive patients with schizophrenia: preliminary results. Schizophrenia Research 50:55–60.

König T, Prichep L, Dierks T, Hubl D, Wahlund LO, John ER, Jelic V (2005) Decreased EEG synchronization in Alzheimer’s disease and mild cognitive impairment. Neurobiology of aging 26:165–171.

Levin R, Nielsen TA (2007) Disturbed dreaming, posttraumatic stress disorder, and affect distress: A review and neurocognitive model. Psychological Bulletin 133:482–528.

Li W, Ma L, Yang G, Gan W-B (2017) REM sleep selectively prunes and maintains new synapses in development and learning. Nat Neurosci 20:427–437.

Makeig S, Inlow M (1993) Lapses in alertness: coherence of fluctuations in performance and EEG spectrum. Electroencephalogr Clin Neurophysiol 86:23–35.

Maris E, Oostenveld R (2007) Nonparametric statistical testing of EEG-and MEG-data. Journal of neuroscience methods 164:177–190.

Marzano C, Ferrara M, Curcio G, De Gennaro L (2010) The effects of sleep deprivation in humans: topographical electroencephalogram changes in non-rapid eye movement (NREM) sleep versus REM sleep. J Sleep Res 19:260–268.

Massimini M, Ferrarelli F, Murphy MJ, Huber R, Riedner BA, Casarotto S, Tononi G (2010) Cortical reactivity and effective connectivity during REM sleep in humans. Cognitive Neuroscience 1:176–183.

Miyauchi S, Misaki M, Kan S, Fukunaga T, Koike T (2009) Human brain activity time-locked to rapid eye movements during REM sleep. Experimental brain research 192:657–667.

Molinari S, Foulkes D (1969) Tonic and Phasic Events during Sleep: Psychological Correlates and Implications. Perceptual and Motor Skills 29:343–368.

Muthuraman M, Hellriegel H, Hoogenboom N, Anwar AR, Mideksa KG, Krause H, Schnitzler A, Deuschl G, Raethjen J (2014) Beamformer Source Analysis and Connectivity on Concurrent EEG and MEG Data during Voluntary Movements. PLOS ONE 9:e91441.

Muzur A, Pace-Schott EF, Hobson JA (2002) The prefrontal cortex in sleep. Trends in cognitive sciences 6:475–481.

Oostenveld R, Fries P, Maris E, Schoffelen J-M (2011) FieldTrip: open source software for advanced analysis of MEG, EEG, and invasive electrophysiological data. Computational intelligence and neuroscience 2011:1.

Oostenveld R, Stegeman DF, Praamstra P, van Oosterom A (2003) Brain symmetry and topographic analysis of lateralized event-related potentials. Clinical Neurophysiology 114:1194–1202.

Perrin F, Pernier J, Bertrand O, Echallier JF (1989) Spherical splines for scalp potential and current density mapping. Electroencephalogr Clin Neurophysiol 72:184–187.

Pruim R, Kaplan DT, Horton NJ (2017) The mosaic Package: Helping Students to’Think with Data’Using R. R Journal 9.

Radua J, Phillips ML, Russell T, Lawrence N, Marshall N, Kalidindi S, El-Hage W, McDonald C, Giampietro V, Brammer MJ, David AS, Surguladze SA (2010) Neural response to specific components of fearful faces in healthy and schizophrenic adults. Neuroimage 49:939–946.

Romero S, Mananas MA, Clos S, Gimenez S, Barbanoj MJ (2003) Reduction of EEG artifacts by ICA in different sleep stages. In: Proceedings of the 25th Annual International Conference of the IEEE Engineering in Medicine and Biology Society (IEEE Cat. No.03CH37439), pp 2675–2678 Vol.3.

Sadaghiani S, Scheeringa R, Lehongre K, Morillon B, Giraud A-L, Kleinschmidt A (2010) Intrinsic connectivity networks, alpha oscillations, and tonic alertness: a simultaneous electroencephalography/functional magnetic resonance imaging study. J Neurosci 30:10243–10250.

Sallinen M, Kaartinen J, Lyytinen H (1996) Processing of auditory stimuli during tonic and phasic periods of REM sleep as revealed by event-related brain potentials. J Sleep Res 5:220–228.

Siegel JM (2005) REM sleep. Principles and practice of sleep medicine 4:120–135.

Siegel JM (2011) REM sleep: a biological and psychological paradox. Sleep Med Rev 15:139–142.

Simor P, Gombos F, Blaskovich B, Bódizs R (2018) Long-range alpha and beta and short-range gamma EEG synchronization distinguishes phasic and tonic REM periods. Sleep 41(3), zsx210.

Simor P, Gombos F, Szakadát S, Sándor P, Bódizs R (2016) EEG spectral power in phasic and tonic REM sleep: different patterns in young adults and children. J Sleep Res 25:269–277.

Spoormaker VI, Gvozdanovic GA, Sämann PG, Czisch M (2014) Ventromedial prefrontal cortex activity and rapid eye movement sleep are associated with subsequent fear expression in human subjects. Exp Brain Res 232:1547–1554.

Srinivasan R, Winter WR, Ding J, Nunez PL (2007) EEG and MEG coherence: measures of functional connectivity at distinct spatial scales of neocortical dynamics. J Neurosci Methods 166:41–52.

Stam CJ, Nolte G, Daffertshofer A (2007) Phase lag index: assessment of functional connectivity from multi channel EEG and MEG with diminished bias from common sources. Hum Brain Mapp 28:1178–1193.

Steriade M, McCormick DA, Sejnowski TJ (1993) Thalamocortical oscillations in the sleeping and aroused brain. Science 262:679–679.

Stuart K, Conduit R (2009) Auditory inhibition of rapid eye movements and dream recall from REM sleep. Sleep 32:399–408.

Tan X, Campbell IG, Feinberg I (2001) A simple method for computer quantification of stage REM eye movement potentials. Psychophysiology 38:512–516.

Tandonnet C, Burle B, Hasbroucq T, Vidal F (2005) Spatial enhancement of EEG traces by surface Laplacian estimation: comparison between local and global methods. Clinical Neurophysiology 116:18–24.

Tarokh L, Carskadon MA, Achermann P (2010) Developmental changes in brain connectivity assessed using the sleep EEG. Neuroscience 171:622–634.

Tarokh L, Van Reen E, LeBourgeois M, Seifer R, Carskadon MA (2011) Sleep EEG provides evidence that cortical changes persist into late adolescence. Sleep 34:1385–1393.

Tenke CE, Kayser J (2012) Generator localization by current source density (CSD): Implications of volume conduction and field closure at intracranial and scalp resolutions. Clin Neurophysiol 123:2328–2345.

Tononi G, Cirelli C (2006) Sleep function and synaptic homeostasis. Sleep medicine reviews 10:49–62.

Trujillo LT, Peterson MA, Kaszniak AW, Allen JJB (2005) EEG phase synchrony differences across visual perception conditions may depend on recording and analysis methods. Clinical Neurophysiology 116:172–189.

Tzourio-Mazoyer N, Landeau B, Papathanassiou D, Crivello F, Etard O, Delcroix N, Mazoyer B, Joliot M (2002) Automated Anatomical Labeling of Activations in SPM Using a Macroscopic Anatomical Parcellation of the MNI MRI Single-Subject Brain. Neuroimage 15:273–289.

Vinck M, Oostenveld R, Van Wingerden M, Battaglia F, Pennartz CM (2011) An improved index of phase-synchronization for electrophysiological data in the presence of volume-conduction, noise and sample-size bias. Neuroimage 55:1548–1565.

Vyazovskiy VV, Olcese U, Hanlon EC, Nir Y, Cirelli C, Tononi G (2011) Local sleep in awake rats. Nature 472:443.

Walker MP, van der Helm E (2009) Overnight therapy? The role of sleep in emotional brain processing. Psychol Bull 135:731–748.

Warrier C, Wong P, Penhune V, Zatorre R, Parrish T, Abrams D, Kraus N (2009) Relating structure to function: Heschl’s gyrus and acoustic processing. J Neurosci 29:61–69.

Waterman null, Elton null, Hofman null, Woestenburg null, Kok null (1993) EEG spectral power analysis of phasic and tonic REM sleep in young and older male subjects. J Sleep Res 2:21–27.

Waterman D, Woestenburg JC, Elton M, Hofman W, Kok A (1992) Removal of ocular artifacts from the REM sleep EEG. Sleep 15:371–375.

Wehrle R, Kaufmann C, Wetter TC, Holsboer F, Auer DP, Pollmächer T, Czisch M (2007) Functional microstates within human REM sleep: first evidence from fMRI of a thalamocortical network specific for phasic REM periods. Eur J Neurosci 25:863–871.

